# *Transmembrane aminopeptidase Q* (*Taqpep*) is a common mechanism in the establishment of periodic patterning in skin and intestine

**DOI:** 10.64898/2026.06.30.735461

**Authors:** Lori B. Dershowitz, Kelly A. McGowan, Ziwei Liu, Brynn M. Brady, Shaul Druckmann, Ulrika Marklund, Gregory S. Barsh, Julia A. Kaltschmidt

## Abstract

Periodic patterns are a frequent motif in biology that occurs across diverse tissues and species. In mammals, pigmentation patterns such as zebra stripes or tiger stripes are well-known examples of periodic patterns; more recently, the myenteric plexus (MP) of the enteric nervous system (ENS), which controls gastrointestinal motility, has been found to exhibit a striped organization in humans and laboratory mice. In domestic cats and other felids, the *Transmembrane aminopeptidase Q* (*Taqpep*) gene plays a key role in color pattern establishment during skin development, but its patterning role has not been examined in other tissues. Here, we show that, in laboratory mice, *Taqpep* is required for normal patterning of developing hair follicles and the MP. Using both sequencing and histologic techniques, we found *Taqpep* is expressed in mesenchymal cells in embryonic skin and intestine directly adjacent to where periodic patterning occurs. We generated *Taqpep* mutant mice, which exhibit disrupted epidermal patterning akin to the changes in periodic coat patterning observed in *Taqpep* mutant cats. The intestine of *Taqpep* mutants has irregularly periodicity of enteric neuronal stripes, and enteric neurons in *Taqpep* mutants exhibit disrupted Wnt signaling. This work provides new insight into the mechanism of enteric neuronal patterning and identify *Taqpep* as a common and conserved mediator of periodic patterning across mammalian tissues and organisms.

**Author summary:** Periodic patterning is a frequent motif in biology. Examples include pigmentation patterning such as tiger stripes and, as recently identified in both mouse and human, the striped organization of enteric neurons in the myenteric plexus of the intestine. In domestic and wild cats, the *Transmembrane aminopeptidase Q* (*Taqpep*) gene is essential for the establishment of periodic patterning. Whether this gene plays a conserved role in periodic patterning across other tissues and species has yet to be explored. We found that *Taqpep* is expressed in mesenchymal cells in embryonic mouse skin and intestine at key locations and developmental stages to instruct periodic patterning. We next generated *Taqpep* mutant mice that exhibit disrupted periodic patterns in both developing skin follicles and in enteric neuron organization. Thus, *Taqpep* is essential in establishing periodic patterning in diverse mammals and tissues.

## Introduction

Periodic patterns are a feature of many tissues across species from cuticle development in *Drosophila* larva to mammalian coat patterning in zebras and cats (1,2). Recently, periodic striping was identified in the mouse and human enteric nervous system (ENS), the branch of the peripheral nervous system that resides in the gut (3,4). After migrating from the neural crest, enteric neurons in the myenteric plexus (MP) become organized into a stripe-like pattern; a developmental event thought to enable the earliest manifestations of gut motility in mid-gestation. We are applying molecular, genomic, and genetic approaches to better understand the mechanisms that underlie ENS patterning, and we hypothesized if genes implicated in other types of periodic patterns might also control the organization of enteric neuronal stripes.

Genetic studies of color patterns in domestic cats have identified a Turing-like process in the embryonic epidermis in which a molecular pre-pattern of long-range Wnt inhibitors and short-range Wnt activators foreshadows the later appearance and form of dark and light markings. A key gene in the establishment of the pre-pattern is *Transmembrane aminopeptidase Q* (*Taqpep*), which encodes transmembrane aminopeptidase Q, a zinc-dependent exopeptidase that is highly conserved across all mammals. The human homolog of *Taqpep* was originally named *Laeverin* based on a high level of expression in chorionic villi. However, several studies including gene-editing in laboratory mice failed to identify a role for the gene in placental development or function (5). Instead, natural mutations of *Taqpep* have been identified as the cause of color pattern abnormalities in domestic cats, cheetahs, and tigers (2,6). In all three species, loss-of-function *Taqpep* alleles cause markings to become more irregular and variable in size and spacing (2,6).

Here we investigate the expression and function of *Taqpep* in the development of hair follicle patterns and intestinal neuronal striping in laboratory mice. We find that *Taqpep* is expressed in embryonic skin and intestine in mesenchymal cells that lie underneath the areas where patterning occurs. Mice homozygous for a gene-edited loss-of-function allele exhibit expanded and disorganized areas of early hair follicle placodes, analogous to what has previously been observed for color pattern in cats. Similarly, enteric neuronal stripes are disorganized, and enteric neurons exhibit a disruption in Wnt signaling. We conclude that *Taqpep* represents a common and conserved molecular signal necessary for normal periodic patterning across species and tissues.

## Results

### Hair follicle patterning is disrupted in *Taqpep* mutant mice

Domestic cats with a *Taqpep* mutation have irregular and expanded dark markings, commonly described as a blotched or marble phenotype. The position and shape of the dark markings are established early in skin development by Wnt activation in thickened areas of the embryonic epidermis and can be visualized by expression of *Dickkopf4* (*Dkk4*), a Wnt target gene that encodes a long-range Wnt inhibitor. Color pattern establishment occurs prior to initiation of hair follicle development which itself is marked by Wnt activation and expression of *Dkk4* in epidermal placodes (7,8).

In mice, periodic patterning of hair follicles begins at embryonic day (E) 13 and is shaped by interactions between the mesenchyme and overlying epithelia (8,9). To assess whether *Taqpep* is positioned temporally and spatially to regulate hair follicle patterning, we used *in situ* hybridization to characterize its expression pattern. At E13, we detected *Taqpep* transcripts in papillary and reticular dermal cells, and in mesenchymal cells adjacent to developing hair buds at E14 (Fig 1A).

**Fig 1.**
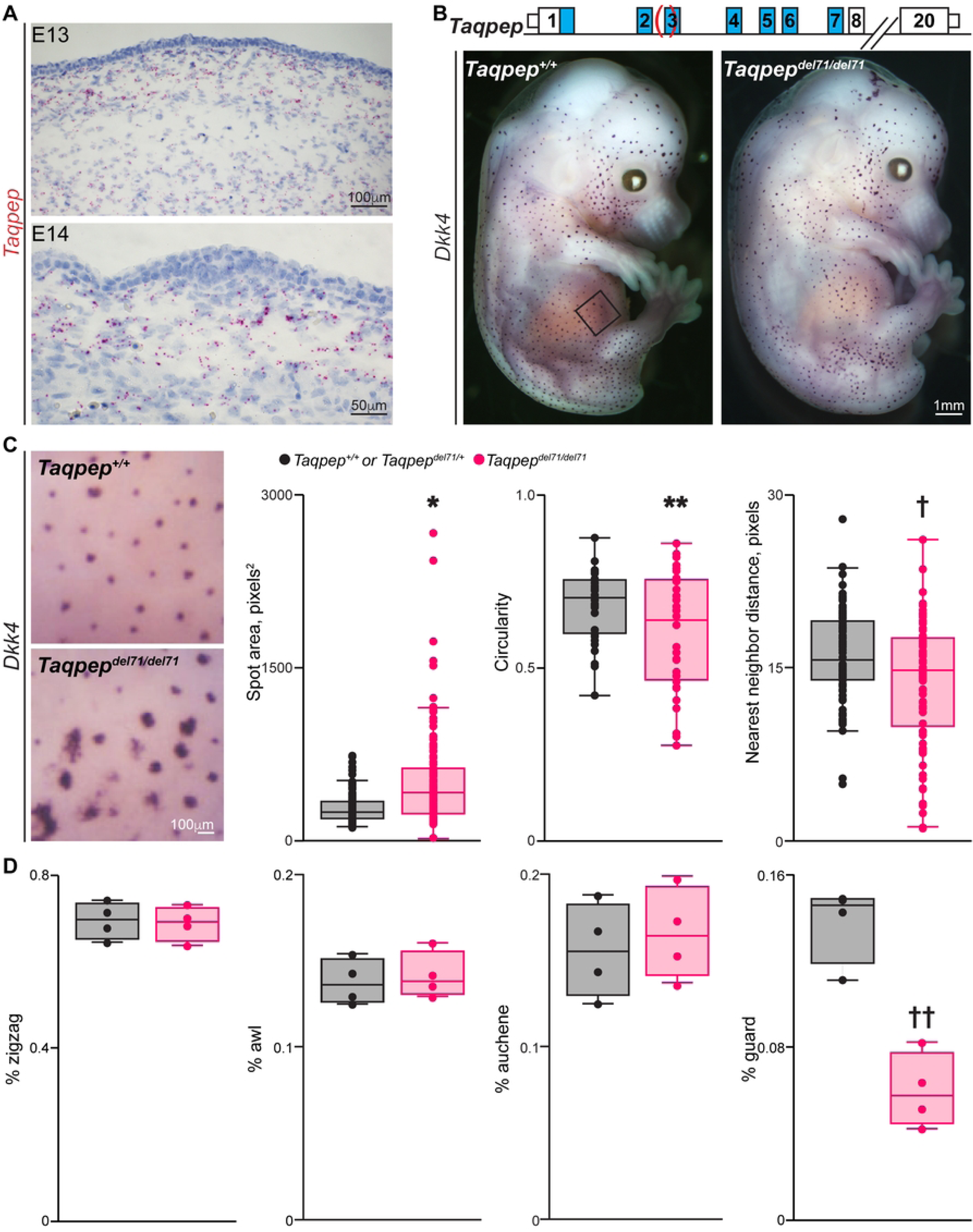
*Taqpep* modifies hair follicle pattern establishment in mice. (**A**) *Taqpep* expression (red puncta) in embryonic mouse dermis (E13) and around a developing hair bud (E14). Photomicrographs are representative images from serially sectioned E13 (n=2 embryos) and E14 (n=1 embryo). (**B, top**). Diagram of mouse *Taqpep* genomic locus (M1 peptidase motif (blue) and del71 mutation (red parentheses, spans 14bp of intron 2 and 57bp of exon 3, chr. 18: 46699451_4699521del, *Mus musculus* GRCm39/mm39)). (**B, bottom**) Wholemount *Dkk4 in situ* hybridization of representative *Taqpep^+/+^* or *Taqpep^del71/^*^+^ and *Taqpep^del71/del71^* E14 embryos identifies early follicle placodes (n=3 embryos of each genotype). Black square denotes anatomic location of images for quantification in c. (**C, left**) Representative images of wholemount *Dkk4 in situ* hybridization from the flank of *Taqpep^+/+^* and *Taqpep^del71/del71^* E14 embryos. Box-whisker plots show spot area, circularity and distance to the single nearest neighbor from *Taqpep^+/+^* or *Taqpep^del71/^*^+^ and *Taqpep^del71/del71^* flank images. Spot area and circularity were evaluated for 152 spots from 3 *Taqpep^+/+^* or *Taqpep^del71/^*^+^ and 121 spots from 3 *Taqpep^del71/del71^* embryos (*p=3.77×10^−8^, **p=0.044, two-tailed t-test). Distance to the nearest neighbor was evaluated for 86 spots from 3 *Taqpep^+/+^* or *Taqpep^del71/^*^+^ and 81 spots from 3 *Taqpep^del71/del71^* embryos (†p=0.0013, two-tailed t-test). The average spot area variance for *Taqpep^+/+^* or *Taqpep^del71/^*^+^ and *Taqpep^del71/del71^* is 15.7+/-3.5 and 155.4+/-57.6, respectively (p=0.036, two-tailed t-test, n=6 photomicrographs for each genotype) and the average circularity variance for *Taqpep^+/+^* or *Taqpep^del71/^*^+^ and *Taqpep^del71/del71^* is 0.0076 +/-0.0009 and 0.0188+/-0.0030, respectively (p=0.012, two-tailed t-test, n=6 photomicrographs for each genotype). The average variance for the five nearest neighbor distances for *Taqpep^+/+^* or *Taqpep^del71/^*^+^ and *Taqpep^del71/del71^* is 34.7 +/- 3.3 and 47.41 +/- 3.8, respectively (p=0.031, two-tailed t-test, n=6 photomicrographs for each genotype). (**D**) Percent of each hair type from the flank of *Taqpep^+/+^* or *Taqpep^del71/+^* and *Taqpep^del71/del71^* adult animals (n=4 animals for each class). ††p=0.0008, two-tailed t-test. Scale bars as indicated.

To explore the role of *Taqpep* in hair follicle patterning, we used CRISPR/Cas9 genome editing and generated a 71-base pair deletion in the *Taqpep* gene in mice. The deletion encompasses the intron 2-exon 3 splice junction (Fig 1B, top) and is predicted to be a loss-of-function allele (See methods, Generation and characterization of *Taqpep^del71^*). We examined expression of *Dkk4* by whole mount *in situ* hybridization in control *Taqpep^+/+^*, *Taqpep^del71/+^* and *Taqpep^del71/del71^* mouse embryos at E14. We found that the regular size and spacing of early follicles observed in *Taqpep^+/+^* and *Taqpep^del71/+^* embryos is disrupted in *Taqpep^del71/del71^* embryos: the dorsal flank lacks an obvious pattern and *Dkk4*+ spots in the ventral flank and thigh are larger, less circular and closer together (Fig 1B, C and Fig S1). Notably, variability in spot area, circularity and distance to the nearest neighbor is greater in *Taqpep^del71/del71^* embryos than in *Taqpep^+/+^*and *Taqpep^del71/+^* embryos (Fig 1C).

We also examined the effect of *Taqpep* on the appearance and distribution of the different adult hair types: zigzag, awl, auchene or guard. We examined 1300-1500 flank hairs collected from four animals of each genotype. The overall appearance of each hair type was similar in all three genotypes. However, we observed an ∼2-fold decrease in the number of guard hairs in the flank of *Taqpep^del71/del71^* animals compared to controls. These observations are consistent with the absence of Dkk4+ spots in the flank of mutant embryos at E14, the developmental stage when the first wave of follicle formation occurs and gives rise to guard hairs.

### *Taqpep* is expressed in the developing intestinal serosa

To determine whether *Taqpep* is expressed in the intestine, we interrogated single-cell RNA sequencing (scRNA-seq) data from embryonic mouse at ages relevant to neuronal stripe development. Enteric neuronal stripes are first evident in the mouse small intestine (SI) at E14.5 and in the colon postnatally (10). Analysis of scRNA-seq data from the E16.5 mouse SI and colon revealed that *Taqpep* expression is enriched in mesothelial cells with some expression in fibroblast-like cells (Fig 2B-E)(11). Thus, *Taqpep* is expressed in mesenchymal cells in the mouse intestine at ages relevant to enteric neuronal stripe development (3,4).

**Fig 2.**
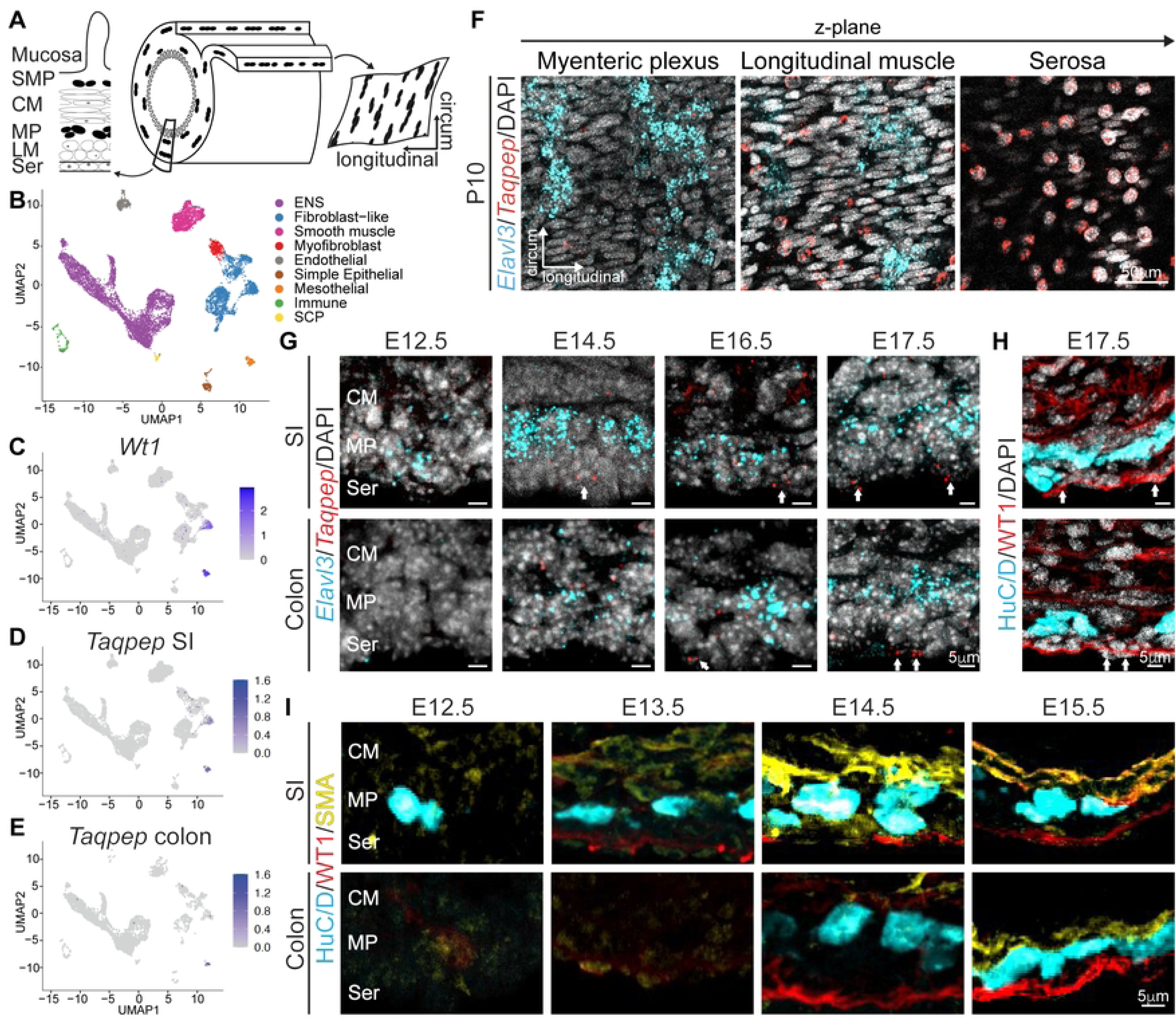
*Taqpep* is expressed in the intestinal serosa. (**A**) Schematic of the intestinal tube with two concentric layers of the enteric nervous system (ENS). On the left, a cross section depicts the intestinal layers. On the right, the muscle and myenteric plexus (MP) layers are peeled to reveal the MP in wholemount. Black dots indicate enteric neurons. (**B**) Uniform manifold approximation and project (UMAP) plot showing single-cell RNA sequencing (scRNA-seq) data from embryonic day (E) 16.5 mouse small intestine (SI) and colon revealing 9 major cell clusters annotated by color. (**C**) Feature plot showing serosal marker *Wt1*. (**D,E**) Feature plots showing *Taqpep* gene expression in subsetted E16.5 SI (D) and colon (E). (**F**) RNAscope with probes against pan-neuronal marker *Elavl3* (cyan) and *Taqpep* (red) through a z-stack from a postnatal day (P)10 control mouse SI. Scale bar as indicated. (**G**) RNAscope with probes against *Elavl3* (cyan) and *Taqpep* (red) in embryonic mouse SI (top) and colon (bottom) from ages spanning E12.5-E16.5. White arrows denote *Taqpep*+ cells. (**H**) Serial sections from the E17.5 SI (top) and colon (bottom) with immunohistochemistry (IHC) labeling for pan-neuronal marker HuC/D (cyan) and serosal marker WT1 (red). White arrows indicate *Taqpep+* WT1+ serosal cells. Scale bars as indicated. (**I**) Representative IHC labeling with HuC/D (cyan), WT1 (red), and smooth muscle marker SMA (yellow) on cross-sections of the embryonic mouse SI (top) and colon (bottom) from ages E12.5-15.5. Scale bars as indicated. CM, circular muscle; LM, longitudinal muscle; MP, myenteric plexus; Ser, serosa; SMP, submucosal plexus.

To visualize *Taqpep* expression in the developing intestine, we performed RNAscope with probes against *Taqpep* and pan-neuronal marker *Elavl3* in cross-sections and MP wholemounts, the layer of the ENS that is striped and controls gastrointestinal motility (Fig 2A)(12). Plane-by-plane analysis at postnatal day (P)10 demonstrates that *Taqpep* is selectively expressed in the serosa (Fig 2F), a monocellular layer of simple squamous epithelium that encases the intestinal tube. Cross-sections across embryonic ages reveal that *Taqpep* is first expressed in the serosa of the SI at E14.5 and of the colon at E16.5 (Fig 2G, arrows). Serial sections of *Taqpep* and Wilm’s tumor protein (WT1), a mesenchymal cell marker that labels serosa, at E17.5 confirm that *Taqpep*+ cells are serosal cells (Fig 2H)(13). WT1+ serosal cells arise in the SI at E13.5 and in the colon at E14.5, approximately 1 day preceding *Taqpep* expression in each location (Fig 2G,H). By E15.5, WT1 labels a continuous outer layer of serosa in both the SI and colon (Fig 2I). To assess the relative position of serosa cells to other cell types throughout embryonic development, we additionally labeled smooth muscle actin and pan-neuronal marker HuC/D. We noticed minimally developed longitudinal muscle as late as E15.5, allowing direct contact between serosal cells and enteric neurons in the embryonic mouse intestine (Fig 2I). Thus, *Taqpep* is expressed in the embryonic intestinal serosa in a spatiotemporal pattern that precedes the emergence of enteric neuronal stripes in the SI and colon (10).

### *Taqpep* mutant mice exhibit altered enteric neuronal stripe organization

We next determined gross intestinal morphology of *Taqpep^del71/del71^*mice. Adult *Taqpep^del71/del71^* mice have no difference in weight, SI length, or colon length as compared to *Taqpep*^+/+^ mice (Fig S2A,C, and G), and *Taqpep* mutant mice generated by other groups also do not demonstrate gross defects (14). Further, hematoxylin and eosin staining of intestine cross sections with accompanying quantification of circular and longitudinal muscle thickness did not reveal any morphological differences between *Taqpep*^+/+^ and *Taqpep^del71/del71^*mice (Fig S2B-I).

To assess whether *Taqpep* influences ENS organization, we examined the position of enteric neurons in MP wholemounts. Circumferentially oriented neuronal stripes were visible in both adult *Taqpep*^+/+^ and *Taqpep^del71/del71^* mice (Fig 3A). The density of HuC/D+ neurons, the area, shape and density of enteric ganglia, and the orientation of neuronal processes are unchanged *Taqpep^del71/del71^* mice (Fig 3B and Fig S3). However, the periodicity of enteric neuronal stripes is altered in *Taqpep^del71/del71^*mice with regions of tightly packed stripes and regions with sparse stripes (Fig 3A). When analyzing distances between the midpoints of adjacent stripes, hereon referred to as “interstripe distance,” we found that while the average interstripe distance remained the same, stripe distribution differed between groups (Fig 3B-F). In *Taqpep*^+/+^ mice, the vast majority of interstripe distances fall within a consistent 200-300 micron range, while in *Taqpep^del71/del71^* mice we observed more occurrences of both, short (<100 microns) and long (>400 micron) interstripe distances (Fig 3D-F). Thus, the regularity of enteric striped patterning is disrupted in *Taqpep* mutant mouse intestine.

**Fig 3.**
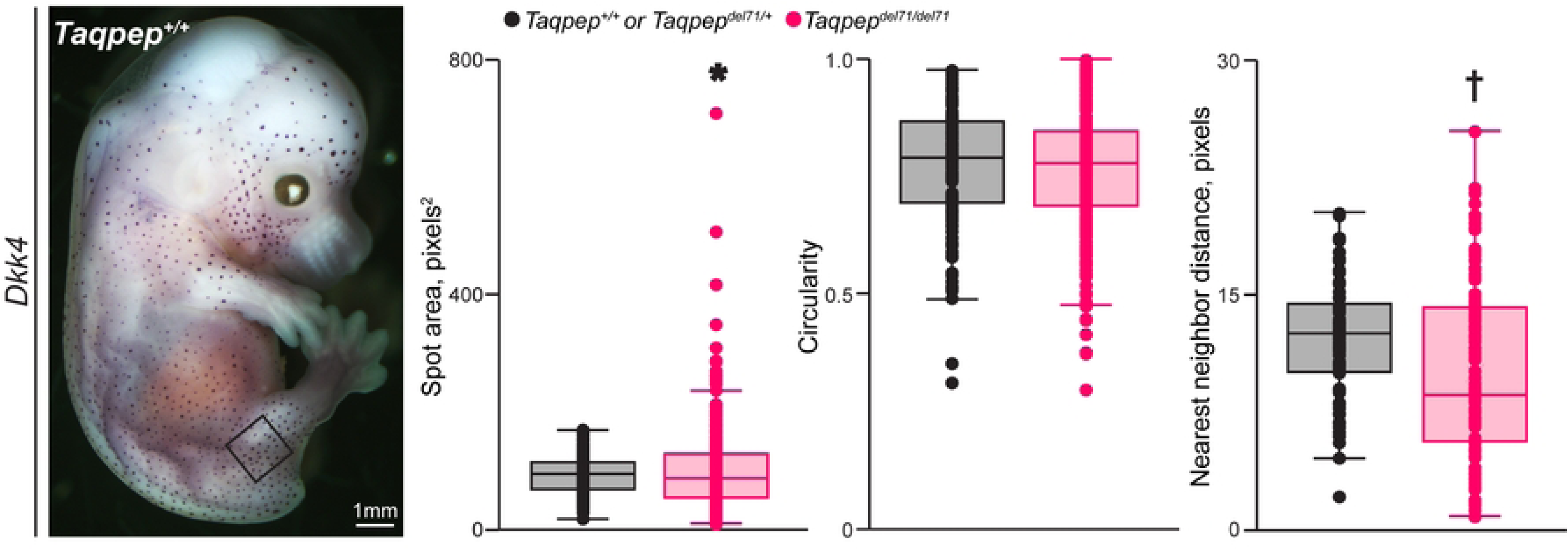
Altered enteric neuronal stripe organization in the *Taqpep^del71/del71^* MP. (**A**) Representative IHC labeling with HuC/D in MP wholemounts from adult *Taqpep^+/+^*(top) and *Taqpep^del71/del71^* ilea (bottom). Interstripe distance is indicated as the distance between the midpoints of serial neuronal stripes. Scale bars as indicated. (**B**) Density of HuC/D+ neurons in MP wholemounts from ilea of adult *Taqpep^+/+^*(black) and *Taqpep^del71/del71^*mice (pink). n=7 *Taqpep^+/+^*, 8 *Taqpep^del71/del71^*. Test unpaired student t-test (mean ± SEM). (**C**) Average interstripe distance derived from IHC labeling with HuC/D in MP wholemounts from adult *Taqpep^+/+^* and *Taqpep^del71/del71^* ilea. n=7 *Taqpep^+/+^*, 8 *Taqpep^del71/del71^*. Test unpaired student t-test (mean ± SEM). (**D**) Standard deviation of interstripe distance from adult *Taqpep^+/+^* and *Taqpep^del71/del71^* ilea. n=7 *Taqpep^+/+^*, 8 *Taqpep^del71/del71^*. Test unpaired student t-test (mean ± SEM). \**p*<0.05. (**E**) Cumulative distribution of interstripe distances from ilea of adult *Taqpep^+/+^* and *Taqpep^del71/del71^* mice. n=7 *Taqpep^+/+^*, 8 *Taqpep^del71/del71^*. Test is bootstrapping of *Taqpep^+/+^* data with 1000 replicates. Reported p is maximum after jackknife resampling. (**F**) Histogram of relative frequency of interstripe distances from adult *Taqpep^+/+^* and *Taqpep^del71/del71^* ilea. n=7 *Taqpep^+/+^*, 8 Taqpep*^del71/del71^*.

### Altered Wnt signaling in the *Taqpep* mutant myenteric plexus

We next sought to determine the downstream mechanism that instructs *Taqpep*-mediated striped patterning in mouse intestine. Altered Wnt signaling modulates hair follicle organization, and Wnt pathway genes establish pre-patterns for skin patterning in both cats and the African striped mouse (6,15–18). Enteric neurons in both mouse and human highly express receptors for Wnt activators and inhibitors, including *Lgr5* and *Lrp5/6* (19,20). To assess for changes in Wnt signaling in the *Taqpep* mutant MP, we intercrossed *Taqpep* mutant mice with *Axin2^LacZ/+^*mice, a reporter line in which cells responding to Wnt signals express β-Galactosidase (βGal)(21). We observed βGal expression in the intestinal crypts, a well-established site of Wnt signaling, as well as in a subset of enteric neurons in the MP (Fig 4A). Co-labeling HuC/D and βGal in MP wholemounts of the early postnatal ileum revealed an increased percentage of MP enteric neurons that express βGal in *Taqpep^del71/del71^* as compared to *Taqpep*^+/+^ mice (Fig 4B,C). Overall, these data suggest that disruption to periodic striped patterning in the *Taqpep* mutant MP may be mediated via the Wnt pathway akin to the observations of disrupted Wnt signaling in *Taqpep* mutant tabby cats (16).

**Fig 4.**
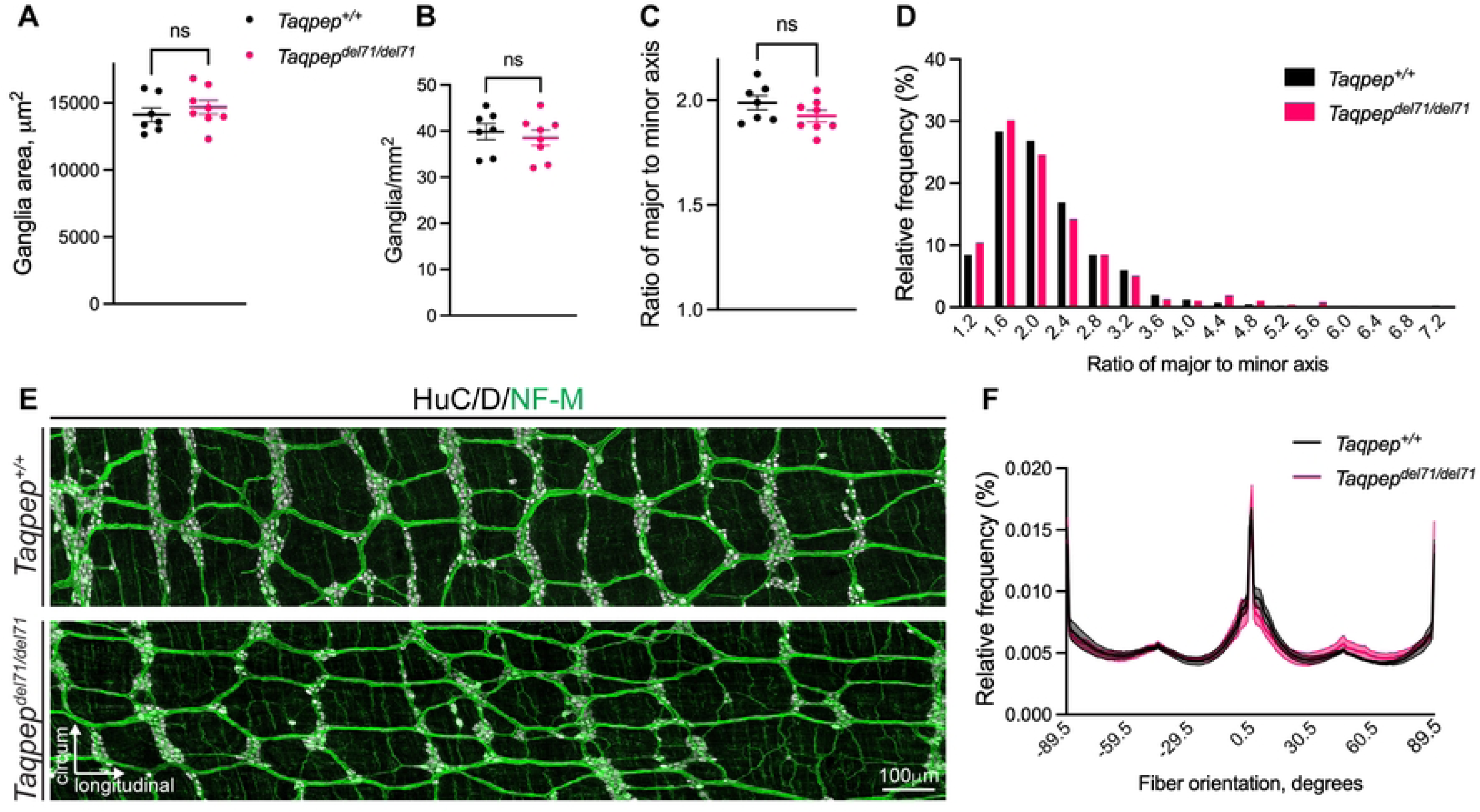
*Axin2^LacZ/+^;Taqpep^del71/del71^* mice have increased Wnt signaling in MP enteric neurons. (**A**) Cross-section of postnatal day (P) 15 *Axin2^LacZ/+^;Taqpep^del71/del71^* ileum with IHC labeling for HuC/D (magenta), β−Galactosidase (β−Gal, green), and DAPI (gray). (**B**) Representative IHC labeling for HuC/D (magenta) and β−Gal (green) in MP wholemounts from P25 ilea in *Axin2^LacZ/+^;Taqpep^+/+^* (left) and *Axin2^LacZ/+^;Taqpep^del71/del71^* mice (right). (**C**) Quantification of percentage of HuC/D+ neurons co-labeled with β−Gal in MP wholemounts of P25 ilea from *Axin2^LacZ/+^;Taqpep^+/+^* (black) and *Axin2^LacZ/+^;Taqpep^del71/del71^* (pink) mice. n= 9 *Taqpep^+/+^*, 8 *Taqpep^del71/del71^*.Test is unpaired student t-test (mean ± SEM); *p = 0.0474. Scale bars as indicated.

## Discussion

The results presented here show that *Taqpep* acts as a periodic patterning gene across different species and organs. Alan Turing first theorized how combinations of diffusible morphogens could generate complex patterns with reaction-diffusion mechanisms (22). Hans Meinhardt and Alfred Gierer refined this model and demonstrated that the combination of long-range inhibitor and short-range activator can generate periodic stripes, and this theory of reaction-diffusion mechanism has been demonstrated experimentally in diverse systems from the striped patterning of the developing mouse hard palate to feather arrangement in developing chick (23–26).

Reaction-diffusion mechanisms involving the Wnt pathway have also been shown to impact hair follicle development and organization in mice (15,17), and a similar mechanism has been proposed to control periodic patterning resulting from downstream effects of *Taqpep* (18,27). Our results support this hypothesis and extend the theory to include periodic patterning of the developing ENS. Though the mechanism by which *Taqpep* modulates Wnt signaling across tissues is unknown, *Taqpep* encodes a metalloprotease with both membrane-bound and secreted forms (28); therefore, Taqpep could act to cleave Wnt-associated activator or inhibitor proteins both locally, as in the physical apposition of the embryonic serosa and MP, or at a distance, as in the *Taqpep* expression surrounding developing hair follicles. In the absence of functional Taqpep, the diffusion range of Wnt activators or inhibitors may change, and mathematical modeling has demonstrated that such a change may result in patterning irregularity (29).

The resultant phenotypes of *Taqpep* loss-of-function mutations in skin and intestine are distinct. In the skin we observed an effect on nearest-neighbor distances, a two-dimensional patterning change, while the ENS exhibited altered neuronal stripe periodicity, a one-dimensional patterning change. Simulation of reaction-diffusion mechanisms on a variety of geometries demonstrated that elliptical geometries, as opposed to rectangular geometries, yield striped patterning even as properties of the boundary conditions are modified. Thus, geometric constraints can impose distinct patterning phenotypes from the same underlying reaction-diffusion mechanism. The different phenotypes may thereby arise from the distinct intrinsic geometry of skin, which is planar, and intestine, which is tubular (30,31).

The observed patterning defect in the ENS of *Taqpep^del71/del71^* mice is reminiscent of an emergent understanding of Hirschsprung’s disease, a developmental disorder of the ENS that results in immotility of the distal colon and inability to pass fecal matter. Historically, defects in Hirschsprung’s disease were thought to be limited to aganglionosis of the distal colon; however, growing clinical literature has reported “skip lesions,” euganglionic regions interspersed within aganglionic regions, and these skip lesions have been observed not only in the colon but also in the ileum (32,33). Current genetic animal models of Hirschsprung’s disease have not been shown to recapitulate skip lesions (34). Here, we demonstrate a genetic means of increasing variability of interstripe distances in the ENS, as the resulting large interstripe distances appear as aganglionic regions interspersed in euganglionic regions. Whether altering the periodicity of enteric neuronal stripes impacts the functioning of the gastrointestinal tract remains to be explored.

## Materials and Methods

### Mice

All experiments followed the National Institutes of Health Guidelines for the Care and Use of Laboratory Animals. The Stanford University Administrative Panel on Laboratory Animal Care approved all procedures. Up to 5 adult mice were kept per cage, and mice received food and water *ad libitum*. For dating embryonic age, female mice were visually inspected each morning for a vaginal plug, and plugged females were separated from the male. Embryos were collected from ages spanning E12.5-E18.5.

Timeline of *Taqpep* expression in the embryonic and neonatal intestine were performed on C57BL/6J mice (The Jackson laboratory).

To generate *Taqpep* mutant mice harboring the *del71* mutation, Cas9 protein and a guide RNA targeting mouse *Taqpep* exon 3 (5’-TTGGACTAAAATATCCCCCCAGG-3’) were microinjected into C57BL/6 zygotes to generate *Taqpep^del71^* animals (chr. 18: 46699450_4699520del, *Mus musculus* GRCm39/mm39). Animals were maintained by backcrossing to C57BL/6 and genotyped using PCR (F-gaaggaatggctttgtggac; R-aacaagggcaacaaggtacg). Experiments were conducted on animals in the F4 generation or later. Two RNA species were identified and sequenced from *Taqpep^del71/del71^* skeletal muscle cDNA: (1) a major product that splices from exon 2 to exon 4 and predicts a frameshift and early termination (NM_029008: c983_1122del predicts NP_083284: pGly277AspfsTer33 and (2) a minor product that splices from exon 2 to intron 3 and predicts an indel in the M1 peptidase domain (g.983_1039delinsAAATATCCCCCCAG which predicts NP_083284:pGln278_His295delinsCysIleSerSerArgPheAlaValIleTrpThr).

To assess for differences in Wnt signaling in *Taqpep^del71/del71^*mice, *Taqpep^del71^*^/+^ mice were crossed to *Axin2^LacZ/+^*mice (The Jackson laboratory) (21). To determine which mice carry a *LacZ* allele, tails were clipped at P7 and placed in X-gal (ThermoFisher) and incubated at 40 ℃ for 1 hour.

### Analysis of single-cell RNA sequencing

We re-analyzed single-cell RNA sequencing (scRNA-seq) data generated previously from the gut of an E16.5 CD1 embryo (11). From the collective dataset iE7.5^Early^-aE16.5, the small and large intestine datasets were merged. Normalization and variance stabilization were performed using SCTransform (v0.4.3), regressing out the mitochondrial percentage. The top 3000 highly variable genes excluding the sex-associated genes (*Xist*, *Gm13305*, *Tsix*, *Gm8730*, *Eif2s3y*, *Ddx3y*, *Uty*, *Kdm5d*) were used for principal component analysis (PCA). Batch correction was performed using Harmony Integration on the first 50 principal components. The resulting Harmony embeddings were subsequently used for k-nearest neighbor(k-NN) graph construction, Leiden clustering, and uniform manifold approximation and projection (UMAP) visualization.

Major cell clusters were manually annotated based on differentially expressed genes identified using the Wilcoxon rank-sum test (Seurat FindAllMarkers function with min.pct=0.5 and logfc,threshold=1), together with the expression of canonical marker genes for gut cell types. Smooth muscle = c(“Myh11”, “Acta2”, “Actg2”),

Fibroblast-like = c(“Col1a1”, “Lum”, “Vim”),

Simple Epithelial = c(“Cdh1”, “Epcam”),

Endothelial = c(“Cd36”, “Pecam1”),

Mesothelial = c(“Upk3b”, “Msln”, “Wt1”,”Taqpep”),

ENS = c(“Phox2b”, “Hand2”, “Ret”),

Immune = c(“Lyz2”, “Mrc1”, “Trbc2”, “Cd52”, “Ptprc”),

Myofibroblast = c(“Trpc3”, “Acta2”, “Tagln”),

SCP = c(“Gfra3”,”Mpz”)

### Intestinal tissue dissection and preparation

For embryonic histology experiments, pregnant females were culled with CO_2_ followed by cervical dislocation. Embryos were removed from the uterus and immediately placed in ice-cold PBS. Extraembryonic membranes were removed, and the intestines were dissected from each embryo. The mesentery was carefully cut away so that the intestines could be pinned in a straight line onto Sylgard 170. While pinned in glass dishes, tissue was fixed in 4% PFA while shaking at 4 ℃ for 1.5 hours. Fixation was followed by 3×10 minute washes in ice-cold PBS. Intestines were stored in 30% sucrose overnight while shaking at 4 ℃. The following day, the intestines were cut into subregions (duodenum, jejunum, ileum, proximal colon, and distal colon), laid on bibulous paper (ThermoFisher), and embedded as cross-sections in OCT (Tissue-Tek) before being stored at -80 ℃. Embedded tissues were sectioned into 12 μm sections at -15 ℃ with a cryostat (Leica CM3050 S).

For neonatal and adult histology experiments, MP wholemounts were collected as previously described (10,35). Briefly, either P10, P15 or 8-10 week-old mice were culled and their intestines were removed. The following tissue segments were collected: 2 cm of duodenum just distal to the stomach, 2 cm of ileum just proximal to the cecum, 2 cm of proximal colon just distal to the cecum, and 2 cm of distal colon just proximal to the rectum. A small cut was made in the proximal end of the tissue to mark the orientation. Tissue pieces were flushed with ice cold PBS, opened along the mesenteric border, and pinned mucosa-side down in a glass dish with Sylgard 170. Tissues were fixed for 1.5 hours in ice-cold PBS while shaking at 4 ℃. Tissues were placed mucosa-side down in ice-cold PBS, and the muscularis (longitudinal muscle, MP, and circular muscle) was peeled with forceps under a dissecting microscope.

### Histology

#### RNAscope *in situ* on intestinal tissues

Detection of mRNA transcripts were performed on either cryosections of mouse embryonic intestine or on wholemounts of P10 SI and colon. For wholemounts, tissues were placed on slides and dried for 2 hours in an oven at 40 ℃. Slides with either embryonic cross sections or neonatal wholemounts were subjected to RNAscope performed per the manufacturer’s guidelines (RNAscope Multiplex Fluorescent V2, ACD) with probes against mouse *Taqpep* (ACD) and *Elavl3* (ACD). Slides were treated with DAPI 1:1000 in PBS for 5 minutes at room temperature, rinsed in ddH_2_O, and mounted with Fluoromount-G (Southern Biotech).

#### *In situ* hybridization on embryonic skin

*In situ* hybridization on paraffin-embedded sections (Fig 1) from 4% paraformaldehyde-fixed samples was performed using the manual RNAscope 2.5 HD Assay-Red Kit (Advanced Cell Diagnostics) according to manufacturer’s instructions. *Taqpep* probes targeted 498-1477 from NM_029008. Images were captured on a Leica DMRXA2 microscope using a Leica DFC550 digital camera and LASV4 (v.4.2.0) software.

For whole-mount *in situ* hybridization, a digoxigenin-labeled RNA probe was generated from a 377bp region spanning exons 3–4 of mouse *Dkk4* using a PCR-based template (cDNA from E12 mouse embryo; AACCACTAAATGGCCAGCAG, ACACAGAATCAATCCCTGGC) and *in vitro* transcription (Roche Diagnostics). Samples were treated with Proteinase K (Sigma), neutralized with 2 mg/ml glycine, fixed in 4% paraformaldehyde with 0.1% glutaraldehyde (Electron Microscopy Sciences), treated with 0.1% sodium borohydride (Sigma), and hybridized with 0.5 μg/ml *Dkk4* riboprobe at 60 °C overnight. Embryos were subsequently treated with 2% blocking reagent (Roche Diagnostics) in maleic acid buffer with Tween-20 (MABT), incubated with an alkaline phosphatase-conjugated anti-digoxigenin antibody (Roche Diagnostics 112093274910, 1:5000) overnight at 4 °C, and developed 3–6 h in BM Purple (Sigma).

#### Immunohistochemistry

For IHC of cryosections, sections were outlined with a hydrophobic barrier pen and washed 3×5 minutes in ice-cold PBS. Primary antibodies were diluted in PBT (PBS, 1% BSA, 0.1% Triton X-100), and sections were incubated in primary antibody solution overnight at 4 ℃ (Table 1).

**Table 1.** Primary antibodies for immunohistochemistry.

| Antibody | Host | Dilution | Source | Catalogue No. | RRID |
| --- | --- | --- | --- | --- | --- |
| βGalactosidase | Rabbit | 1:5000 | MP Bio | 02104939-CF | AB_2335286 |
| Calretinin | Rabbit | 1:2000 | Millipore | AB5054 | AB_2068506 |
| HuC/D | Human | 1:128,000 | Gift of Vanda Lennon | A-21272 | AB_2813895 |
| Neurofilament-M | Rabbit | 1:2000 | Millipore | AB1987 | AB_91201 |
| nNOS | Sheep | 1:1000 | Millipore | AB1529 | AB_90743 |
| Somatostatin | Rat | 1:500 | Millipore | MAB354 | AB_2255365 |
| Smooth muscle actin | Goat | 1:500 | Abcam | AB21027 | AB_1951138 |
| WT1 | Mouse | 1:500 | Santa Cruz Biotechnology | sc-393498 | AB_2905496 |

The following day, slides were washed 3×5 minutes in ice-cold PBS before adding secondary antibodies diluted in PBT (Table 2) at room temperature for 2 hours. Slides were washed 3×5 minutes in room temperature PBS and rinsed with ddH_2_O before mounting with Fluoromount-G (Southern Biotech).

**Table 2.**
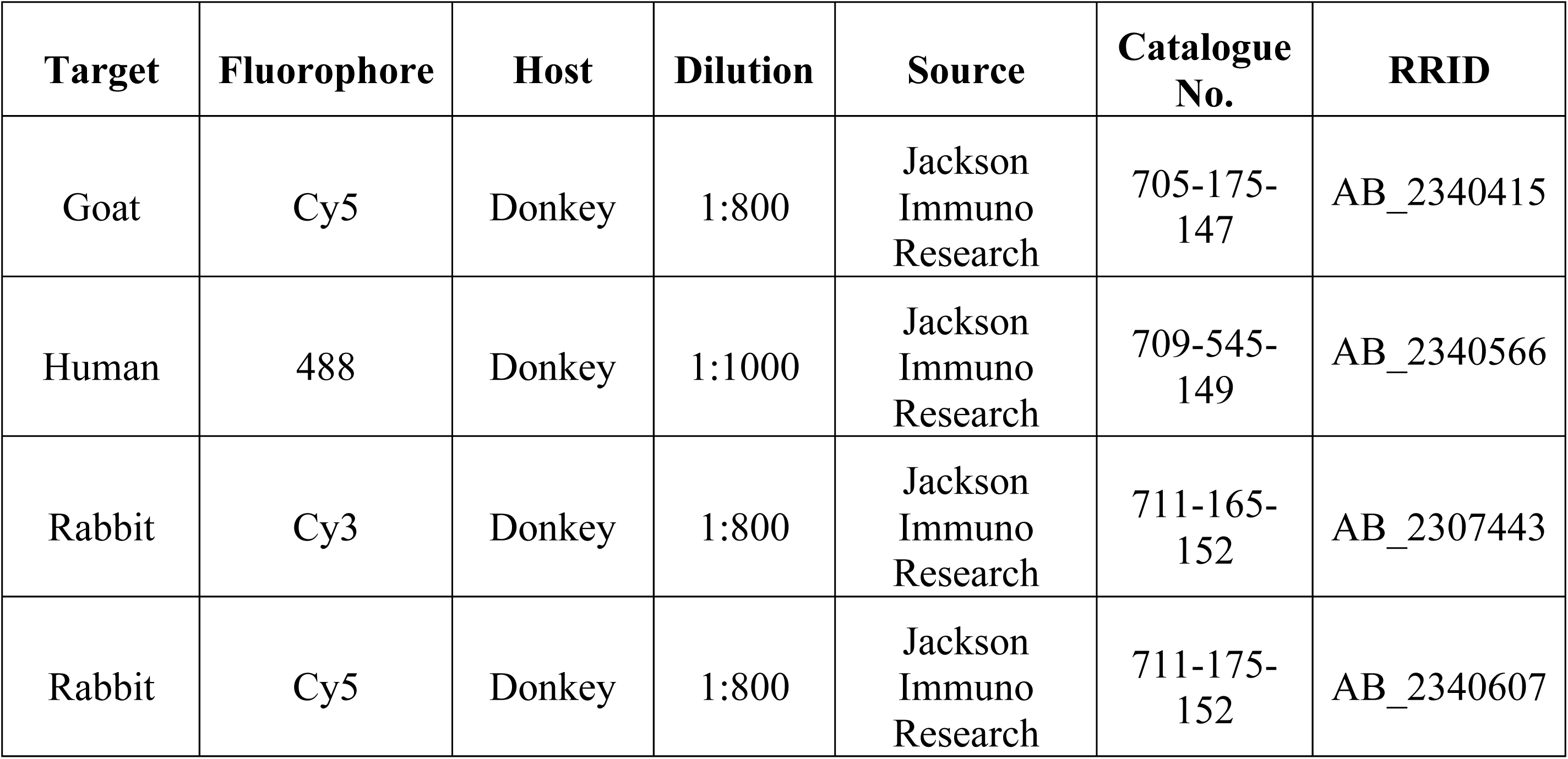
Secondary antibodies for immunohistochemistry.

| Target | Fluorophore | Host | Dilution | Source | Catalogue No. | RRID |
| --- | --- | --- | --- | --- | --- | --- |
| Goat | Cy5 | Donkey | 1:800 | Jackson Immuno Research | 705-175-147 | AB_2340415 |
| Human | 488 | Donkey | 1:1000 | Jackson Immuno Research | 709-545-149 | AB_2340566 |
| Rabbit | Cy3 | Donkey | 1:800 | Jackson Immuno Research | 711-165-152 | AB_2307443 |
| Rabbit | Cy5 | Donkey | 1:800 | Jackson Immuno Research | 711-175-152 | AB_2340607 |

IHC of wholemounts were performed as previously described (10). Briefly, 2 cm pieces of muscularis tissue were transferred to WHO microtitration trays (International Scientific Supplies) with PBS. Tissue was transferred to primary antibody diluted in PBT (Table 1) and incubated overnight while shaking at 4 ℃. The next morning, tissue was washed 3×30 minutes in PBT at room temperature before transferring to secondary antibody diluted in PBT (Table 2) for 2 hours at room temperature. Tissue was washed 2×10 minutes in PBT, 2×10 minutes in PBS, and 1×5 minutes in DAPI 1:1000 in PBS. Tissue was mounted on slides with a paintbrush, rinsed in ddH_2_O, and mounted with Fluoromount-G (Southern Biotech).

### Image acquisition

All fluorescent microscopy images were acquired with a 20x (NA 0.75) oil objective on a Leica SP8 confocal microscope. For wholemount images, regions of interest were selected, imaged, and stitched with Navigator mode in LASX (Leica) as previously described (10). All images were acquired as z-stacks with either 2 μm between planes for embryonic ages or 3 μm between planes for postnatal ages.

Skin whole-mount preparations were photographed on a dissecting microscope under direct illumination.

### Image quantification

Image analyses were performed in ImageJ/FIJI (NIH, Bethesda, MD). Neuronal density of HuC/D maximum intensity projections was performed as previously described (10).

Quantification of HuC/D and enteric neuronal subtype marker co-labeling was performed as previously described (10).

### Patterning analysis

#### Skin patterning

Spot pattern was evaluated from a 1.25 mm x 1.25 mm square that lies halfway between the axillary and inguinal crease (flank), or a square centered on the upper thigh. Spot area, circularity and nearest neighbor distance were evaluated using Fiji. Circularity is spot area/(π x major radius^2^). For the nearest neighbor analysis, the distance to the five closest spots was measured for all spots in which the 5 nearest neighbors are closer than the spot is to the photograph edge.

#### ENS patterning

To analyze interstripe distances, a 500 μm tall region that spanned the length of the tissue from proximal to distal was selected from HuC/D maximum intensity projections of 8-12 week-old *Taqpep^+/+^*and *Taqpep^del71/del71^* mice. Images were blurred and thresholded in FIJI. Images were blinded, and a grid with 100×100 mm regions was projected onto the image with the grid tool in FIJI. Along each horizontal line of the grid (5 lines per image spaced 100 μm apart), the line segment tool was used to mark the midpoint of each neuronal stripe intersected, and the Measure Segmented Distances in ImageJ macro was used to measure the distances between midpoints of neuronal stripes (10).

To analyze ganglia area, density, and the ratio of major to minor axes, the same blurred HuC/D maximum intensity projections were used as above. Due to intrinsic differences in HuC/D intensity across the length of an image, subsections of each image were thresholded and binarized in FIJI and restitched in Adobe Photoshop. To identify individual ganglia and approximate each as an ellipse, the measure region properties function was utilized in scikit-image (36). To avoid biasing the data toward small groups of cells rather than large clusters, ganglia < 500 μm^2^ in diameter were manually excluded.

To assess the orientation of Neurofilament-M+ neuronal fibers, microscopy images of Neurofilament-M in wholemounts from *Taqpep^+/+^*and *Taqpep^del71/del71^* mice were blurred and thresholded. OrientationJ was used to determine the representation of neuronal fibers at different orientations with 0° defined as horizontal (proximal-distal) (37).

### Hair type quantification

Hair was plucked from the flank (3cm from the hind limb on the right side) of 2-3-month-old *Taqpep^+/+^* and *Taqpep^del71/del71^* animals (n=4 animals for each genotype). 1300-1500 hairs from each animal were classified according to hair type.

### Figures, graphing and statistical analysis

Figures and graphics were constructed in Adobe Illustrator 2024. For image analysis, all cropping and rotations were performed in ImageJ. For fluorescent microscopy images included in figures, levels of individual channels were linearly changed in Adobe Photoshop 2024 with all changes applied equally between images of *Taqpep^+/+^* and *Taqpep^del71/del71^* mice.

All graphing and statistical analyses were performed with Prism 9 software (GraphPad). Unpaired student t-tests were used to compare the means of data quantified from *Taqpep^+/+^* and *Taqpep^del71/del71^* mice. Bootstrapping of *Taqpep^+/+^* interstripe distance data with 1000 replicates was compared to *Taqpep^del71/del71^* mutant distributions to determine the significance of the difference between *Taqpep^+/+^*and *Taqpep^del71/del71^* cumulative distribution functions. For this analysis, significance was reported as maximum p-value after jackknife resampling.

## Data availability

All correspondence should be directed to Dr. Gregory Barsh at and Dr. Julia Kaltschmidt at. All data and code are available upon request to the corresponding author after publication. At the time of review, all data and code are available to reviewers upon request.

## Acknowledgments

We would like to thank all members of the Kaltschmidt lab for feedback on experimental design. The HuC/D primary antibody was a generous gift from Vanda Lennon at Mayo Clinic. Funding acknowledgements include: Stanford Medical Scientist Training Program grants T32 GM007365-44, T32 GM145402 (LBD); Swedish Research Council (Vetenskapsrådet; 2020-01129 and 2022-01570) (UM); European Research Council (divENSify; 101045026) (UM); Knut and Alice Wallenberg Foundation (KAW; 2020.0109) (UM); Strategic Research Area in Stem Cell Research and Regenerative Medicine (UM); National Institutes of Health grants AR067925 and AR082708 (KAM, GSB); National Institutes of health grant R21 HD110950 (JAK); Firmenich Foundation (JAK); Wu Tsai Neurosciences Institute (JAK); Stanford University Department of Neurosurgery (JAK).

## Author contributions

LBD contributed to conceptualization, methodology, investigation, funding acquisition, and writing. KAM contributed to conceptualization, methodology, investigation, and writing. ZL contributed to analysis. BMB contributed to investigation. SD contributed to methodology. UM contributed to methodology, analysis, and conceptualization. GSB contributed to conceptualization, methodology, funding acquisition, writing, and supervision. JAK contributed to conceptualization, methodology, investigation, funding acquisition, writing, and supervision.

## Figure legends

**Fig S1.**
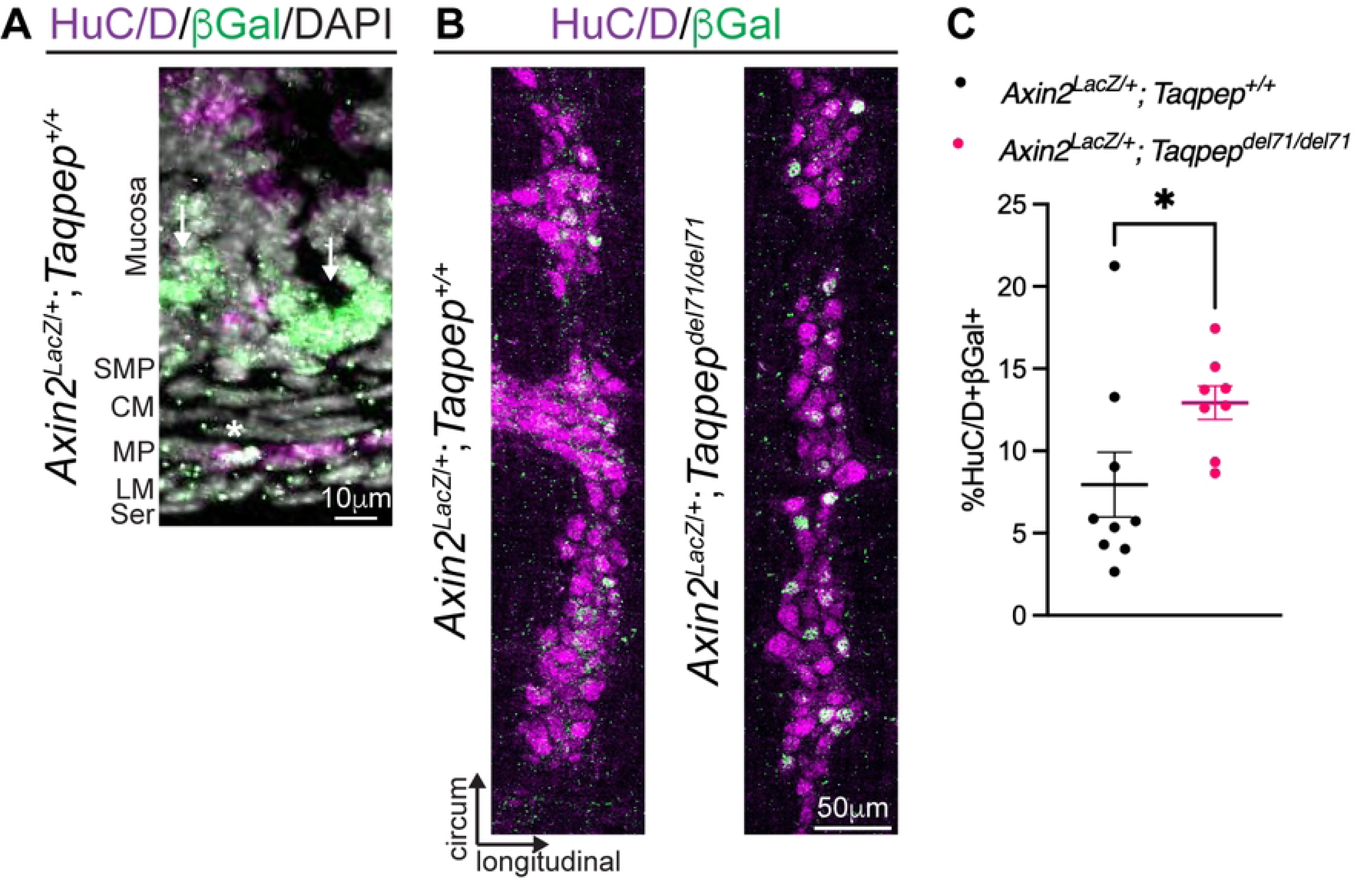
Thigh hair follicle patterning in *Taqpep^del71/del71^* embryos. Black square denotes anatomic location of images used for quantification. Box-whisker plots show spot area, circularity and distance to the single nearest neighbor from *Taqpep^+/+^* or *Taqpep^del71/^*^+^ and *Taqpep^del71/del71^* thigh images. Spot area and circularity were evaluated for 139 spots from n=3 *Taqpep^+/+^* or *Taqpep^del71/^*^+^ and 197 spots from n=3 *Taqpep^del71/del71^* embryos (*p=0.025, two-tailed t-test). Distance to the nearest neighbor was evaluated for 78 spots from n=3 *Taqpep^+/+^* or *Taqpep^del71/^*^+^ and 118 spots from n=3 *Taqpep^del71/del71^* embryos (†p=0.0013, two-tailed t-test). The average spot area variance for *Taqpep^+/+^* or *Taqpep^del71/^*^+^ and *Taqpep^del71/del71^* is 944 +/- 159 and 6861 +/- 1015, respectively (p=0.027, two-tailed t-test, n=4 *Taqpep^+/+^* or *Taqpep^del71/^*^+^ and n=6 *Taqpep^del71/del71^* photomicrographs) and the average variance for the five nearest neighbor distances for *Taqpep^+/+^* or *Taqpep^del71/^*^+^ and *Taqpep^del71/del71^* is 29.3 +/- 2.8 and 41.8 +/- 4.0, respectively (p=0.035, two-tailed t-test, n=4 *Taqpep^+/+^* or *Taqpep^del71/^*^+^ and n=6 *Taqpep^del71/del71^* photomicrographs). Scale bar as indicated.

**Fig S2.**
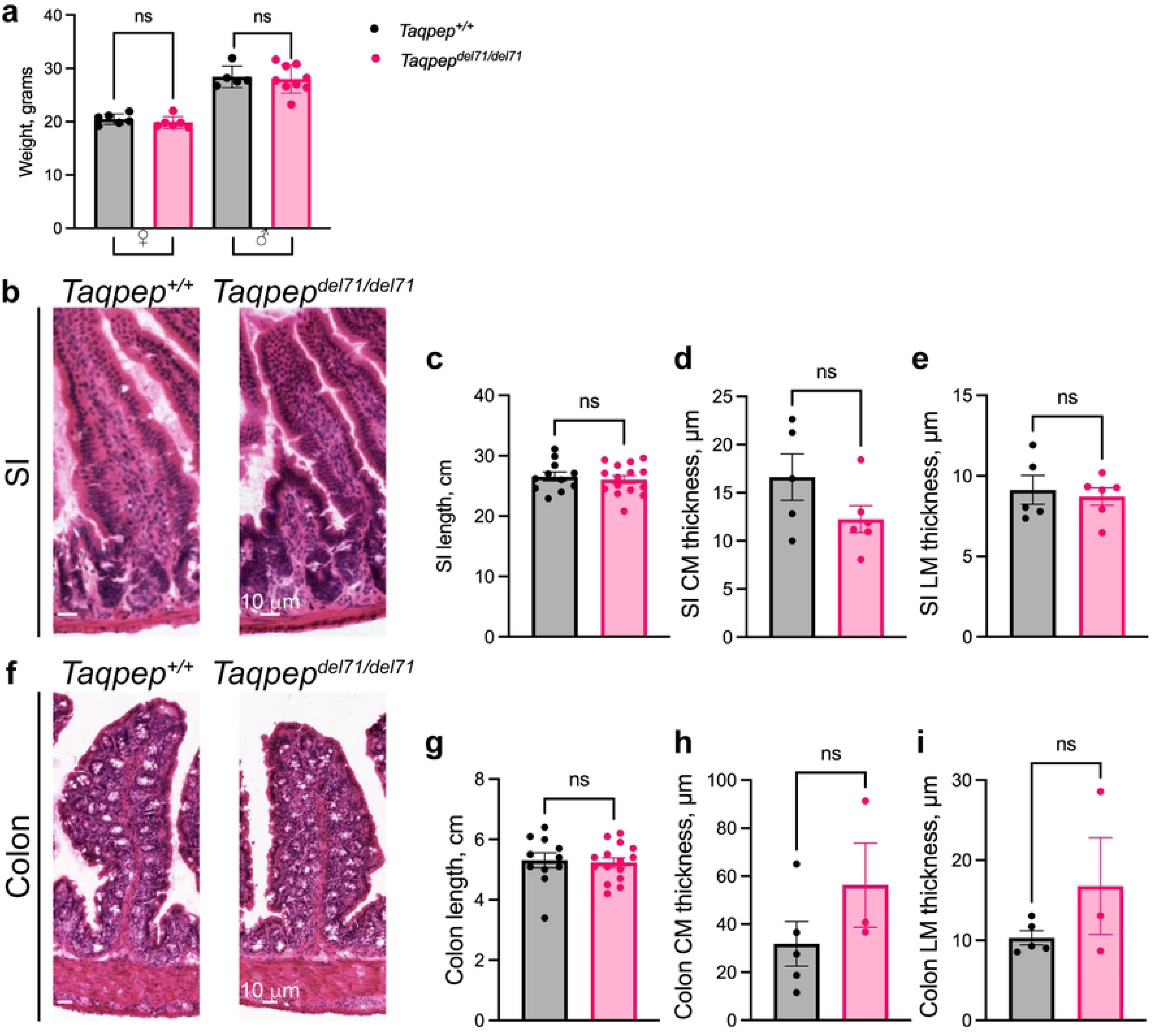
*Taqpep^del71/del71^* mice do not exhibit gross gastrointestinal defects. (**A**) Weight of adult female (left) and male (right) *Taqpep^+/+^* and *Taqpep^del71/del71^* mice. (**B**) Hematoxylin and eosin staining of cross-sections from the small intestine (SI) of adult *Taqpep^+/+^* (left) and *Taqpep^del71/del71^* mice (right). (**C**) Quantification of SI length in adult *Taqpep^+/+^* and *Taqpep^del71/del71^* mice. (**D, E**) Quantification of circular muscle thickness (D) and longitudinal muscle thickness (E) derived from hematoxylin and eosin staining of SI cross-sections in adult *Taqpep^+/+^* and *Taqpep^del71/del71^* mice. (**F**) Hematoxylin and eosin staining of cross-sections from the colon of adult *Taqpep^+/+^* (left) and *Taqpep^del71/del71^* mice (right). (**G**) Quantification of colon length in adult *Taqpep^+/+^* and *Taqpep^del71/del71^* mice. (**H,I**) Quantification of circular muscle thickness (H) and longitudinal muscle thickness (I) derived from hematoxylin and eosin staining of colon cross-sections in adult *Taqpep^+/+^* and *Taqpep^del71/del71^* mice. All tests unpaired student t-tests (mean ± SEM). Scale bars as indicated.

**Fig S3. Enteric ganglia and neuronal fibers are unaffected in *Taqpep^del71/del71^* mice.**

(**A,B**) Quantification of the density (a) and area (b) of ganglia in *Taqpep^+/+^* (black) and *Taqpep^del71/del71^* ilea (pink). n=7 *Taqpep^+/+^*, 8 *Taqpep^del71/del71^.* All tests unpaired student t-tests (mean ± SEM). (**C,D**) Quantification (C) and distribution (D) of the ratio of the major to minor axis for ganglia approximated as an ellipse from *Taqpep^+/+^* (black) and *Taqpep^del71/del71^* ilea (pink). n=7 *Taqpep^+/+^*, 8 *Taqpep^del71/del71I^.* All tests unpaired student t-tests (mean ± SEM). (**E**) Representative IHC labeling with HuC/D (gray) and Neurofilament-M (NF-M, green) in wholemounts of the ileum MP from adult *Taqpep^+/+^* (top) and *Taqpep^del71/del71^* mice (bottom). (**F**) Quantification of relative frequency of NF-M fiber orientations observed in the ileum MP from *Taqpep^+/+^* (black) and *Taqpep^del71/del71^* mice (pink). A fiber orientation of 0 is defined as horizontal. n=6 *Taqpep^+/+^*, 5 *Taqpep^del71/del71^*. Scale bar as indicated.

